# Whole-cell particle-based digital twin simulations from 4D lattice light-sheet microscopy data

**DOI:** 10.1101/2025.04.09.647865

**Authors:** Eric Arkfeld, Zichen Wang, Hiroyuki Hakozaki, Johannes Schöneberg

## Abstract

We present a framework for performing whole-cell digital twin simulations which integrates 4D (x,y,z,t) lattice light-sheet microscopy (LLSM) data with particle-based reaction-diffusion modeling to capture intracellular organelle dynamics. Using imaging data from Cal27 cells, we construct digital twins that incorporate mitochondrial networks, microtubule networks, dynein and kinesin motors, the plasma membrane, and the nucleus. Passive diffusive mitochondrial dynamics are parameterized using stochastic reaction-diffusion simulations in ReaDDy, while active transport is modeled explicitly by incorporating motor-driven transport along a diffusing, polarized microtubule network. Our simulations accurately reproduce experimentally observed mitochondrial dynamics across pharmacological microtubule depolymerization conditions and reproduce the mitochondrial response to intermediate perturbations without explicit re-parameterization. This novel meso-scale digital twin framework offers a bridge between atomic-scale whole-cell simulations and experimental time and length scales.

## INTRODUCTION

Whole-cell models (WCMs) aim to provide a predictive in-silico framework for understanding cellular processes by integrating diverse molecular, structural, and biophysical data into computational simulations. Bridging molecular and cellular scales, WCMs can help elucidate complex biological processes such as gene regulation, metabolism, and phenotypic variability at the cellular level^1,2^. Owing to such unique insights, the applications of WCMs range from drug discovery and personalized medicine, where they could be used to model disease states and predict drug responses, to synthetic biology and metabolic engineering, where they could facilitate the rational design of genetic circuits and improve efficiency in industrial biotechnology.

Historically, WCM development has been tightly coupled to advancements in experimental techniques as their construction and parameterization are extremely data intensive (as recently reviewed here^3,4^). Early WCMs relied on quantitative metabolic assays alongside numerical flux balance analysis (FBA) to predict cellular growth and metabolic states for *Escherichia coli*. Enabled by advances in multi-omics, Karr, et al. constructed a WCM for *Mycoplasma genitalium* which modeled metabolism, transcriptional regulation, RNA processing, protein synthesis, and DNA replication using a hybrid computational framework^5^. Early 3D (x, y, z) WCMs introduced explicit spatial effects via static excluded volumes representative of organelles^6^. In their recent 3D-WCM of JCVI-syn3A^7^, Thornburg et al. used structural data obtained from cryo-electron tomography (CET) to define the spatial distribution of ribosomes and membrane geometry.

Despite such advances in WCMs, current approaches remain limited. Mesoscale intracellular processes such as organelle dynamics, spatial exclusion, and complex confinement geometries all modulate local biochemical environments on timescales of seconds. These effects can be captured through explicit particle-based reaction-diffusion simulations where several high-performance computational frameworks have been developed to address this need^8–15^. We hypothesized that there exists an intermediate mesoscopic level of detail where, with the correct approximations, realistic simulations of cell biology may be performed using such tools. Live cell microscopy data of organelle dynamics would be ideal for constructing and parameterizing such simulations. However, the high phototoxicity of common live-cell fluorescence microscopy techniques (e.g., confocal) has so far hindered the development of such mesoscale 3D-WCM simulations^4^. With current bottom-up 3D-WCMs (e.g. coarse-grained molecular dynamics (CG-MD), genome-scale models) being constructed from static structural data (e.g., CET), the incorporation of such spatiotemporal dynamics remains challenging.

Lattice light-sheet microscopy (LLSM) is an advanced four-dimensional (4D; x, y, z, t) imaging technique which enables high-speed, high-resolution volumetric data acquisition of cellular and subcellular structures with minimal photodamage^16,17^. LLSM illuminates samples with a thin sheet of structured light, maintaining a spatial resolution of ∼250 nm laterally and ∼500 nm axially with a frame rate of up to 1 second per volumetric frame. This makes LLSM particularly suitable for long-term imaging of subcellular dynamics in live cells. Leveraging this 4D imaging data, recent advances in image preprocessing^18^ and image analysis^19^ have enabled quantification of organelle dynamics such as mitochondria.

Beyond their classic role as the powerhouse of the cell, mitochondria are implicated in signaling, calcium homeostasis, and cell fate decisions^20–22^. Mitochondrial dysfunctions are implicated in a range of diseases ranging from rare primary mitochondrial disorders to common conditions like diabetes, neurodegeneration, and cancer^23–27^. In mammalian cells, mitochondrial number, volume fraction, and morphology vary by cell type and state. Exhibiting morphologies ranging from small vesicular puncta to extensive tubular networks, mitochondrial networks are continuously remodeled through fusion and fission^28^. Fusion—the merging of multiple mitochondria into a single entity—facilitates material exchange, enhances reactant uptake for ATP production, and buffers damaged mitochondrial DNA, reactive oxygen species, and cytosolic calcium^20,29^. Conversely, fission—the fragmentation of a mitochondrion into multiple units—isolates damaged regions for degradation, enables active transport of fragments along the cytoskeleton, and partitions mitochondria during cell division^30,31^. These remodeling events are tightly regulated and responsive to changes in energetic state, environmental stressors, and drug-induced perturbations^31^. Due to their critical role in cell biology and their rich behavior and dynamics, mitochondria are an excellent model system for the development of mesoscale WCMs.

Simulations have been successfully used to gain insights into, and make predictions about the structure, dynamics, and quality control mechanisms of mitochondrial networks. While prior works have modeled mitochondria at different scales using a variety of approaches^32–35^, only recently has reaction-diffusion dynamics been used to explicitly model mitochondrial dynamics^36^. However, existing mitochondria models remain fundamentally limited as they fail to account for a core conserved process exhibited by mitochondria in eukaryotic cells: active transport along a microtubule network.

Here we present the first-in-class mesoscale digital twin whole-cell modeling and simulation framework that reliably simulates the mitochondrial network, including the microtubule network, molecular motors, and cellular membranes. Using live Cal27 cells as the model system, each whole-cell digital twin model incorporates 3D particle-based reconstructions of the mitochondrial network, microtubule network, nuclear membrane, and plasma membrane constructed from the imaging data of its biological counterpart. By creating an automated model construction pipeline, we allow for rapid and reproducible construction of digital twin models directly from standard segmentation outputs. We used ReaDDy^14,15^ to develop and perform all simulations. Building complexity in a stepwise manner, we established a baseline by simulating passive mitochondrial dynamics in Cal27 cells treated with nocodazole to disrupt the microtubule network. Leveraging the high spatial and high temporal resolution afforded by LLSM, passive models were parameterized using the same dataset used for model construction. Next, we developed an active transport module for ReaDDy which enabled the first motor-driven active transport simulations in somatic cells. We utilized this to simulate mitochondrial dynamics with motor-driven active transport along an explicitly diffusing, polarized microtubule network model reconstructed from imaging data of untreated control cells. To validate our approach, we tested whether our model could predict the effects of partial disruption of the microtubule network without reparameterization. Using the parameter set for the control condition, the simulated digital twin models successfully predicted intermediate trends in mitochondrial dynamics with a partially disrupted microtubule network. Our framework provides a powerful platform for investigating whole-cell dynamics, bridging the mesoscale gap to enable the simulation and prediction of cellular behavior within a biologically realistic context.

## RESULTS

### Creating a whole-cell digital twin for simulating passive mitochondrial network dynamics

Our first objective was to create a particle-based digital twin model to serve as the initial state for passive mitochondrial network dynamics simulations. We used Cal27 cells, a head and neck squamous carcinoma cell line^37^, as the biological model system. Mitochondria are known to diffuse or be actively transported along both microtubule and actin networks, where microtubule-mediated are dominant^38^. To focus on building a working passive diffusion model, active transport contributions to mitochondria motility were suppressed by treating the cells with nocodazole for 60 minutes to disrupt the microtubule networks^19,39^. 4D fluorescence microscopy data was collected for live Cal27 cells using our custom-built LLSM at a rate of 11 seconds per volumetric frame (171µm x 96µm x 22µm) for a total of 60 frames (11 minutes). MitoTracker Green was used to fluorescently label mitochondria and Tubulin Tracker Deep Red was used to fluorescently label microtubules. A lack of fluorescent signal in the deep red channel confirmed disruption of the microtubule network.

The first step in creating a mesoscale digital twin WCM is to create initial coordinates for the simulation. After preprocessing the raw imaging data (**Methods**), cells and their nuclei were segmented in 3D using the first frame of the 4D LLSM imaging data according to the spatial exclusion of mitochondrial fragments (**Figure 1A**). This was then used to create the whole-cell digital twin model (**Figure 1B**). Specifically, each 3D digital twin model (**Figure 1C**) includes particle-based representations for the cell’s plasma membrane (red), the nuclear membrane (blue), and the mitochondrial network (green). Mitochondrial fluorescence was segmented in each volume using MitoGraph^40^ (**Figure 1D**). The discretized MitoGraph segmentation outputs (**Figure 1E**) were then used to generate the particle-based reconstruction of the cell’s mitochondrial network (**Figure 1F**). Next, we set out to obtain the parameters that would govern the particle-based reaction diffusion simulation. The spatiotemporal evolution of the mitochondrial network (**Figure 1G**) was quantitatively tracked using MitoTNT^19^ (**Figure 1H**). Briefly, MitoTNT tracks the mitochondrial network by utilizing distance and topology continuity at high frame rates which allows for the quantification of mitochondrial motilities and fusion/fission rates.

**Figure 1.**
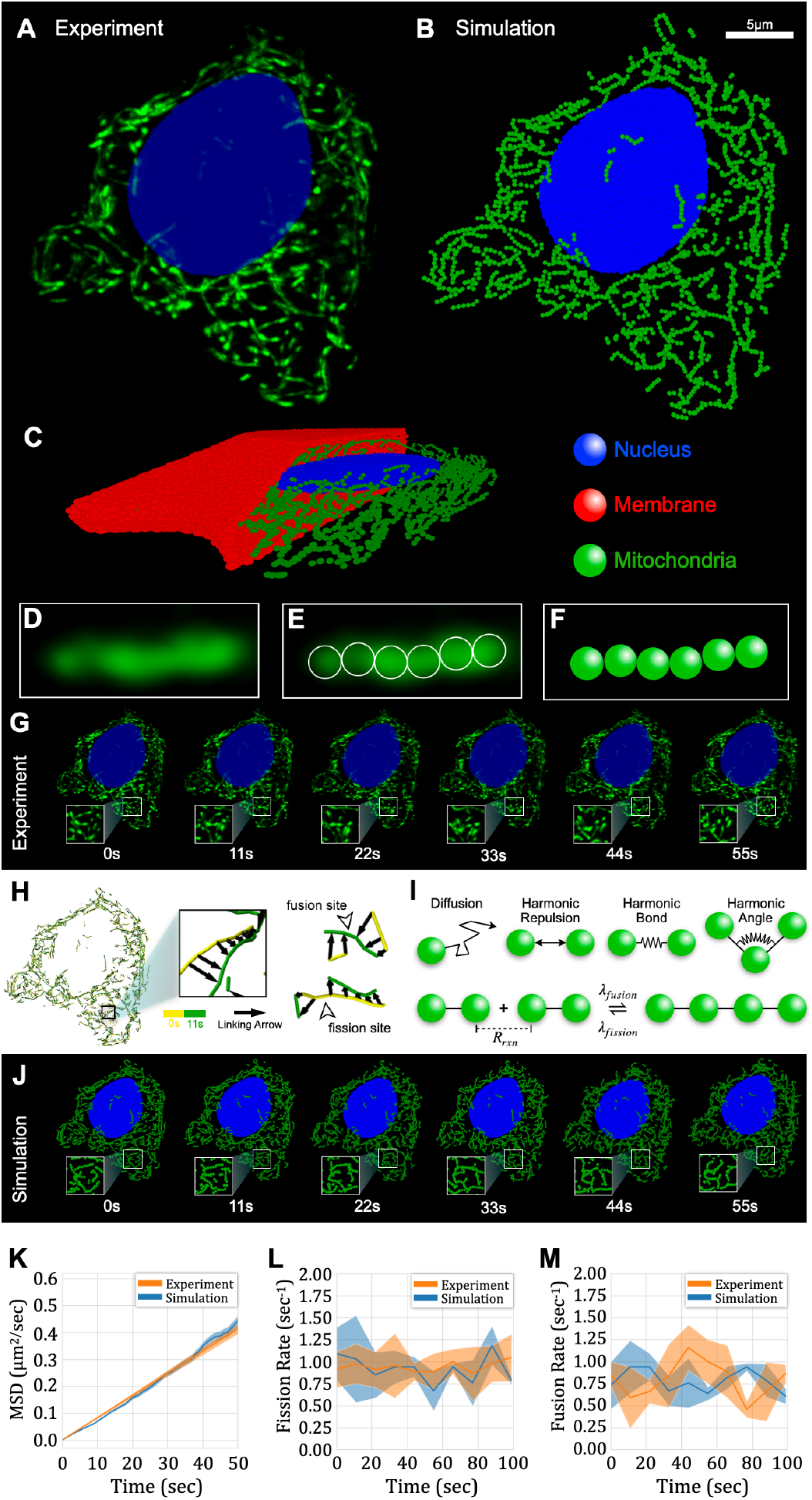
Whole-cell digital twins can be constructed that accurately simulate passive mitochondrial dynamics. (A) First frame of 4D LLSM data of a single Cal27 cell treated with nocodazole for 60 minutes and labeled with MitoTracker (mitochondria, green). The nucleus (blue) and membrane (hidden for visibility) were reconstructed based on the spatial exclusion of mitochondria. Cells were additionally strained with Tubulin Tracker Deep Red where the lack of fluorescence confirmed disruption of the microtubule network. (B) In-silico particle-based reconstruction of the same Cal27 cell including the mitochondrial network (green), nucleus (blue) and membrane (red, hidden for visibility). Simulations are rendered using Simularium Viewer^42^. (C) Model side view with partially hidden membrane (red) which envelopes the mobile species and defines the simulation domain. (D) Example mitochondria fluorescence signal segmented using MitoGraph^40^ (E) The particle-based model is constructed from using the resulting network graph structure by replacing nodes with particles at the specified coordinate while the cell’s membrane and nucleus were constructed from segmentation masks by placing particles along the mask boundary. (F) Mitochondria particles are given a radius of 0.15 µm, are assigned 3D coordinates. (G) Representative 4D LLSM imaging data collected with a frame interval of 11 seconds. (H) MitoTNT was used for tracking mitochondrial fragments across successive frames and for quantifying motility and remodeling events. (I) Particle interactions and reactions implemented in the passive twin simulation including diffusion, harmonic potentials, and topology reactions. (J) Representative frames from a simulated trajectory for the digital twin illustrating mitochondrial network dynamics. Frames were sampled every 11 seconds to match the experimental frame interval. (K) Simulated and experimental fragment-level MSD versus time for calibrated microscopic diffusion coefficients. Shaded regions correspond to the standard-error of the mean. (L) Simulated and experimental macroscopic fission rates versus time. Shaded regions correspond to the standard deviation. (M) Simulated and experimental macroscopic fusion rates versus time. Shaded regions correspond to the standard deviation. See Simularium link in text to view a trajectory for this cell.

We constructed the digital twin models using ReaDDy^14,15^. Using the mitochondrial network graphs obtained from image segmentation, particle-based representations of the mitochondrial network were generated by mapping nodes and edges to discrete particles and harmonic bonds (**Figure 1I**). Mitochondria particles were given a fixed radius of 0.15 µm to reflect the typical cross-section radii for mitochondria^41^. An equilibrium bond length of 0.3 µm was used to ensure that bonded particles remained spatially adjacent. We constrained fragment topologies by defining harmonic angle potentials between all sets of three consecutive bonded mitochondria particles. Mitochondrial fission and fusion events were modeled as topological reactions that alter fragment topologies by adding or removing harmonic potentials between particles. Mitochondrial fusion was modeled as a spatially dependent topology reaction which adds a bond between particles with a fixed rate if within a predefined threshold distance. Mitochondrial fission was modeled as structural topology reactions where bonds are randomly removed at a fixed rate at each integration timestep. To restrict the mitochondrial network dynamics to the cytosolic compartment while reproducing geometric constraints, we created 3D particle-based reconstructions of each cell’s nuclear and plasma membranes using their respective segmentation masks (**Methods**). The positions of nuclear and plasma membrane particles were fixed as minimal movement was observed in the LLSM imaging data over the duration of imaging (**Figure S1**). Both species were assigned a radius of 0.2 µm as we found it provided sufficient structural detail in both cases without excessive particle counts that lead to increased computational cost. Spatial exclusion was imposed using piecewise repulsive harmonic potentials between all particle species using the sum of their respective radii as the cutoff distances. The resulting digital twin models provide a well-defined initial state within the current simulation scope. We were able to automate the entire model construction pipeline starting from segmentation outputs. Requiring less than 1 minute on a workstation, this pipeline allows efficient yet consistent construction of digital twin models directly from segmented microscopy data.

### Whole-cell digital twin simulations of passive mitochondrial dynamics

We modeled the time-evolution of the digital twin model using ReaDDy’s overdamped Langevin dynamics (**Figure 1J**) where all simulations were performed using a 5 ms integration timestep and at a temperature of 37°C which matches the temperature of our live-cell imaging. Reactions were evaluated using a Kinetic Monte Carlo approach as implemented in ReaDDy’s Gillespie stochastic simulation algorithm (SSA)^14,43^.

Passive mitochondrial motility was modeled using diffusive motion by tuning the microscopic diffusion coefficient to reproduce the ensemble averaged fragment-level mean-squared displacement (MSD) observed for the Cal27 cell mitochondria treated with nocodazole for 60 minutes (**Figure 1K**). The simulated MSD curve closely matches experimental observations with a comparable standard error of the mean (SEM) (**Figure 1K, shaded region**). Harmonic repulsion and bond potentials were calibrated to maintain fragment topologies still enforcing spatial exclusion for the 5 ms timestep (**Figure S2**). Assuming an equilibrium angle of π radians, we calibrated harmonic angle potentials to recapitulate the observed fragment-level topologies under the wormlike chain model by tuning the force constant to reproduce the mean-squared end-to-end distance for linear mitochondrial fragments (**Methods**).

Fission rates obtained from MitoTNT analyses were directly used as the simulation fission rate parameter. The resulting fission rate versus time plot demonstrates strong agreement between experimental and simulated data, with fluctuations captured within one standard deviation (**Figure 1L**). Due to its bimolecular nature, the microscopic fusion rate constant was iteratively refined to maintain the experimentally observed mean fragment length (**Methods**). The resulting macroscopic fusion rate also aligns well with experimental measurements, exhibiting comparable dynamics and variability as indicated by the standard deviation (**Figure 1M**). These results validate the parameterization approach and confirm that the digital twin model successfully captures key aspects of passive mitochondrial dynamics within live human cells. A trajectory for this cell can be viewed in your browser using the following Simularium link: https://tinyurl.com/Cal27-noco60min

### Creating whole-cell digital twins for simulating mitochondrial network dynamics with active transport along a polarized, diffusing microtubule cytoskeleton

Our next objective was to incorporate an explicit microtubule model to model active mitochondrial transport. 4D imaging data was collected using LLSM for live, untreated Cal27 cells using a frame interval of 11 seconds over the course of 60 frames (11 minutes). Mitochondria and microtubules were fluorescently labeled using MitoTracker Green and Tubulin Tracker Deep Red. The first frame of the 4D imaging data (**Figure 2A**) was used to construct a particle-based whole-cell digital-win model (**Figure 2B**). In addition to the particle-based reconstructions for the mitochondria, plasma membrane, and nuclear membrane, each 3D model (**Figure 2C**) now incorporates an explicit polarized microtubule model with explicit motor particle species (**Figure 2D**).

**Figure 2.**
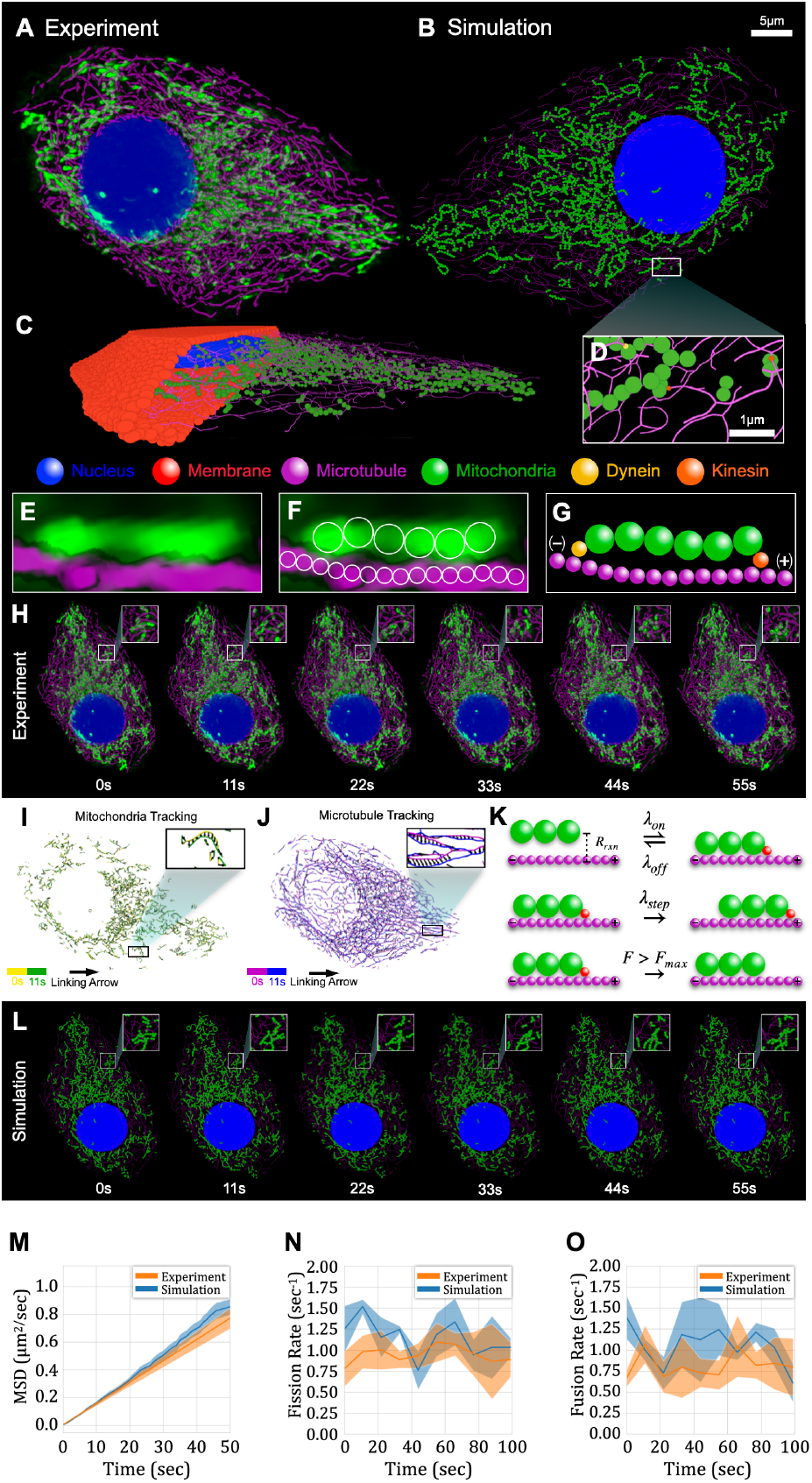
Whole-cell digital twins can be constructed that accurately simulate active mitochondrial dynamics on a diffusing microtubule network. (A) First frame of 4D LLSM data of a control Cal27 cell labeled with MitoTracker (mitochondria, green) and Tubulin Tracker Deep Red (microtubules, magenta). The nucleus (blue) and membrane (hidden for visibility) were reconstructed based on spatial exclusion of the mitochondria. (B) Particle-based digital twin for the same control Cal27 cell where the mitochondrial network (green), microtubule network (magenta), nucleus (blue), and membrane (hidden for visibility) are all constructed from 4D LLSM imaging data. Simulations are rendered using Simularium Viewer^42^. (C) Side view with partially hidden membrane (red) which confines the mobile species to the cytosolic compartment. (D) Zoomed view of the simulation with mitochondria undergoing antero- and retrograde transport by kinesin (orange) and dynein (orange) motor particles walking along the polarized, diffusing microtubule network. (E) Mitochondria fluorescence signal (green) is segmented using MitoGraph and the tubulin fluorescence signal (magenta) was segmented using the Allen Institute Cell Segmentation^44^ plugin for Napari^45^ and was processed using MitoGraph^40^ to obtain data in standard format. (F) Particle-based models were constructed from spatially embedded mitochondria and microtubule network graphs by replacing nodes with particles. (G) Schematic of a simulated mitochondrial fragment (green) bound the polarized passively diffusing microtubule (magenta) via explicit kinesin (orange) and dynein (yellow) motor particles. Note that motor particles are not derived from imaging data but are dynamically added and removed over the course of the simulation according to binding/unbinding reaction events. (H) Representative 4D LLSM data of mitochondria and microtubules collected using a frame interval of 11 seconds. (I) MitoTNT tracking of the mitochondria network dynamics used for parameterization of mitochondrial model dynamics. (J) MitoTNT tracking of the microtubule network dynamics used for parameterization of microtubule model dynamics. (K) Schematic diagram of the reactions for modeling active transport. (L) Representative trajectory frames from the digital twin simulation illustrating both microtubule and mitochondrial network dynamics. Frames were sampled at an interval of 11 seconds to match the experimental frame interval. (M) Simulated and experimental fragment-level MSD versus time for calibrated microscopic diffusion and active transport parameters. Shaded regions correspond to the SEM. (N) Simulated and experimental macroscopic fission reaction rates versus time. Shaded regions correspond to the standard deviation. (O) Simulated and experimental macroscopic fusion rate constant versus time. Shaded regions correspond to the standard deviation. See Simularium link in text to view a trajectory for this cell.

To create the initial particle coordinates of the model, microtubule fluorescence was segmented using the Allen Cell & Structure Segmenter^44^ with default 3D settings for alpha tubulin (**Figure 2E**) in Napari^45^. The resulting binary microtubule mask was then processed using a custom python script to obtain an undirected, spatially embedded graph-based reconstruction of the cell’s microtubule network (**Figure 2F**). Mapping nodes and edges to particles and harmonic bond potentials, we additionally defined distinct antero- and retrograde directions for each microtubule (**Methods**) to enable motor driven transport by kinesin and dynein species (**Figure 2G**). Briefly, we achieved this by first introducing a spatially embedded centrosome node based on the distinct morphology and elevated fluorescence intensity of the centrosome relative to the surrounding microtubules in the imaging data. Next, we connected this centrosome node to proximal nodes in the microtubule network directed all edges in the retrograde direction based on the shortest path to the centrosome node. For any remaining disjoint graphs, a terminal (degree=1) node was randomly selected, and all edges were directed towards it in similar fashion. We extended our automated model construction pipeline to reconstruct the cell’s microtubule network as a spatially embedded directed graph from 4D microscopy data.

To parameterize the dynamics of the model from experimental results, the spatiotemporal dynamics of both the mitochondrial network and microtubule network (**Figure 2H**) were analyzed using MitoTNT. Mitochondrial network dynamics were analyzed as described for the passive model (**Figure 2I**). With the segmented microtubule network sharing similar properties with the mitochondrial network (e.g., distinct local topology, small displacements at high framerates), we used MitoTNT to track and extract morphological and temporal features from the segmented 4D microtubule data (**Figure 2J**).

### Parameterization and Validation of Active Transport along the Microtubule Cytoskeleton

We modeled anterograde and retrograde transport of mitochondria along the polarized, diffusing microtubule network by extending ReaDDy to incorporate a custom active transport module. Each motor particle is treated as an independent realization of a two-state Markov process with active and inactive states. ReaDDy reactions (**Figure 2K**) are used to dynamically add and remove explicit kinesin and dynein motor particle species throughout the course of the simulation **(Figure 2L**). Motors are initially considered inactive, with activation governed by a predefined transition probability. Regardless of state, motor positions are corrected each timestep to remain fixed relative to the diffusing, polarized microtubule. At each timestep, motors may activate, deactivate, step, or detach. Active motors stochastically step in a directed manner along the microtubule. Multiple motors can independently bind to one or more particles within a single topology; Here, we restricted motor binding to terminal mitochondria particles with one motor per site. Fragments may be transported by one or more motors which are in turn bound to one or more microtubules. Motors detach upon reaching filament end or if the bond force with mitochondria exceeds a specified threshold.

The microscopic diffusion coefficient for the microtubule particles was calibrated to reproduce ensembled node-level MSD obtained from MitoTNT analyses. To confine the microtubule network to the cytosolic compartment, microtubule particles were given a radius of 0.05 µm for defining repulsive potentials with the nuclear and plasma membrane particles and harmonic angle potentials were introduced as conformational constraints. We parameterized motor dynamics primarily from experimentally determined microkinetic values obtained from literature sources (**Table S3**). We assumed that all motors bind with the same rate where the average microkinetic binding rate was used globally. Consequently, the only free active transport parameter was the motor activation probability, which we subsequently tuned to reproduce the fragment diffusivities averaged across four untreated (control) Cal27 cells (**Figure 2M**). Except for fusion rates, mitochondria utilized parameters obtained from passive simulations including fission rates (**Figure 2N**). Fusion rates with active transport (**Figure 2O**) were iteratively tuned to maintain constant fragment size. The simulated fission and fusion rates over time closely align with experimental data, capturing fluctuations within the standard deviation. A trajectory for this cell can be viewed in your browser using the following Simularium link: https://tinyurl.com/Cal27-control

### Digital twins can predict the mitochondrial dynamical response to intermediate nocodazole perturbations without explicit parameterization

To assess the capability of our digital twin simulations to predict mitochondrial network responses to intermediate disruption of the microtubule network, we simulated mitochondrial dynamics digital twins constructed for Cal27 cells subjected to 30 minutes of nocodazole treatment and compared the results to experimental observations. With nocodazole inducing progressive depolymerization of the microtubule network as a function of incubation time^39^, we hypothesized that trends in mitochondrial motility and remodeling events would be captured by the digital twin simulations without reparameterization.

Compared to the untreated control cells (**Figure 3A**), LLSM imaging of the data of cells treated for 30 minutes (**Figure 3B**) revealed significant but incomplete disruption of the microtubule network as indicated by the reduction in normalized fluorescence intensity. For the cells treated for 60 minutes, the microtubule signal was absent except for the centrosome (**Figure 3C**). Quantification confirmed that a 30 min nocodazole treatment reduced the microtubule fluorescence intensity to about half that of the untreated condition (**Figure 3D**).

**Figure 3.**
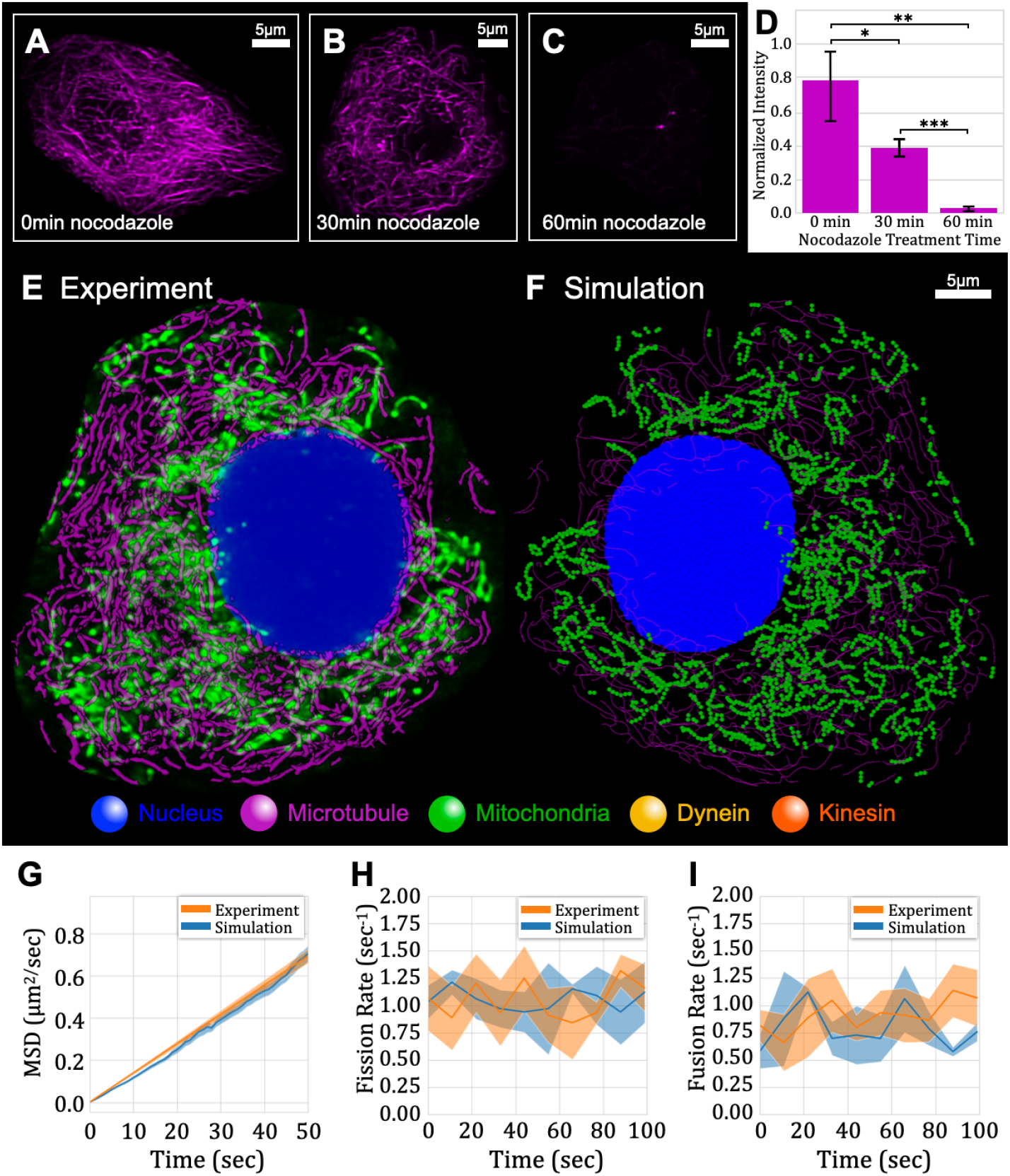
Simulations accurately predict drug-induced perturbations in mitochondrial dynamics caused by microtubule disruption. (A) Microtubule fluorescence intensity for the untreated (0 min) Cal27 control cell shown in Figure 2. (B) Microtubule fluorescence intensity for a Cal27 cell treated with nocodazole for 30 minutes. (C) Microtubule fluorescence intensity for the Cal27 cell treated with nocodazole for 60 minutes shown in Figure 1. (D) Comparison of fluorescence intensities for 3-sets of 4x Cal27 for cells treated with nocodazole for 0 (control), 30, and 60 minutes, respectively. Intensity values are normalized with respect to cytosolic volume and to per-cell intensities across all cells in all conditions. (E) First frame of 4D LLSM imaging data of a Cal27 cell treated with nocodazole for 30 minutes with MitoTracker (mitochondria, green) and Tubulin Tracker Deep Red (microtubules, magenta). (F) The particle-based digital twin for the same Cal27 cell. Model components include the mitochondrial network (green), microtubule network (magenta), nucleus (blue), and membrane (hidden for visibility). Simulations are rendered using Simularium Viewer^42^. (G) Simulated and experimental fragment-level MSD) versus time. Shaded regions correspond to the SEM. (H) Simulated and experimental macroscopic fission reaction rates versus time. Shaded regions correspond to the standard deviation. (I) Simulated and experimental macroscopic fusion rate constant versus time. Shaded regions correspond to the standard deviation. See Simularium link in text to view a trajectory for this cell.

The 4D LLSM dataset for Cal27 cells incubated with nocodazole for 30 minutes (**Figure 3G**) was used to construct digital twin models (**Figure 3H**), which consequently included the partially disrupted microtubule network. Simulations of the time evolution of these digital twins were performed using parameters derived from the untreated control cells. We found that simulated fragment-level MSDs were in excellent agreement with experimental observations (**Figure 3I**). As hypothesized, mitochondria in the 30-minute condition exhibited reduced motility relative to the control but remained higher than that of the 60-minute condition. We additionally found that the digital twin simulations recapitulated both fission (**Figure 3J**) and fusion (**Figure 3K**) event rates observed experimentally for their live counterparts. These results demonstrate that our digital twin simulations can accurately capture mitochondrial responses to pharmacological perturbation to the microtubule network without reparameterization. A trajectory for this cell can be viewed in your browser using the following Simularium link: https://tinyurl.com/Cal27-noco30min

## DISCUSSION

Here we presented a first-in-class framework for constructing and simulating whole-cell digital twins by integrating 4D live-cell microscopy data with ReaDDy’s particle-based reaction-diffusion dynamics for modeling organelle dynamics within cells. LLSM was used to obtain 4D data on mitochondrial and microtubule network dynamics in Cal27 cells across three conditions where the cell’s microtubule network was disrupted to varying degrees. To facilitate model construction, we created an automated pipeline which uses standard segmentation tool outputs to generate whole-cell digital twin models incorporating spatially resolved 3D particle-based reconstructions of the mitochondrial network, microtubule network, plasma membrane, and nuclear membrane. Spatial and topological constraints were implemented using harmonic potentials. Passive transport is modeled using particle-based reaction-diffusion dynamics using a Langevin thermostat at a temperature of 37°C. To model active transport, we developed a custom ReaDDy module which enables explicit motor particles (e.g., kinesin and dynein) which dynamically bind, unbind, activate, and step stochastically along a passively diffusing, polarized microtubule network. Motors additionally exhibit force-dependent behavior where excessive strains result in immediate unbinding from the microtubule which mimics experimentally observed force-dependent detachment. Using simulation parameters derived from literature and MitoTNT analyses performed on the same experimental data, the resulting digital twin simulations reproduce trends in mitochondrial motility and network remodeling rates experimentally observed in their live biological counterparts across all three treatment conditions.

Our mesoscale modeling approach demonstrates that mitochondrial network dynamics can be essentially accurately captured without the necessity for full molecular-level detail. Despite the inherent assumptions required at the mesoscale, our simulations nonetheless reproduce the core features of mitochondrial behaviors and implies that we have sufficient understanding of the underlying physics. This highlights an important opportunity: There exists a relevant mesoscale at which simplified, spatially explicit, and physically principled simulations can both explain observed biological phenomena and generate biologically relevant, experimentally accessible predictions about cellular behavior. This mesoscale level of simulation represents a first-in-class addition to existing approaches such as coarse-grained atomistic models (e.g., Martini^46,47^) or genome-scale models which encode the cell’s reactions in a spatial lattice (e.g., Lattice Microbes^48^). Our framework demonstrates that lightweight, spatially embedded particle-based models offer a powerful and extensible compromise—capturing emergent organelle-level behaviors while remaining modular, computationally tractable, and readily adaptable to new experimental data.

As the first mesoscale digital twin approach to WCM modeling, our current implementation has several immediate limitations. While we incorporate an explicit polarized microtubule network which diffuses, our model does not yet account for dynamic processes such as microtubule polymerization, depolymerization, and nucleation. While our simulations accurately capture mitochondrial dynamics, they do not yet account for interactions with other organelles such as the endoplasmic reticulum which are known to play a crucial role in mitochondrial network regulation. Lastly, although we implemented force-dependent detachment for molecular motor species, our current implementation lacks force-dependent procession, motor-cargo adapters, and microtubule associated proteins (MAPS).

Moving forward, several key developments will enhance the biological realism and predictive capabilities of our digital twin simulations. Incorporating additional organelles such as the ER will improve the representation of organelle-organelle interactions. Improving the molecular detail of active transport with force-dependent procession, motor adaptors, and MAPS would enrich intracellular dynamics. Modeling dynamic cytoskeletal rearrangements and plasma membrane topologies would enable the study of cell-wide morpho dynamics under varying physiological conditions. Hybridizing our current framework and adding a grid-based model for the cytoplasm would enable concentration dependent effects and intracellular signaling events to be modeled. Validation of our whole-cell digital twin simulations across additional cell types, perturbations, and disease models will provide insights into mitochondrial dysfunction in neurodegenerative diseases, cancer, and metabolic disorders. Such extensions and integrations could provide a path for the development of biologically realistic simulations that capture the emergent structure-function relationships characteristic to the basic unit of life: the cell.

## METHODS

### Cell Culture

Cal27 cells were cultured in 90% DMEM, high glucose, GlutaMAX (Gibco, #10566016), 10% FBS (GenClone, #25-550), 1% Penicillin-Streptomycin (Gibco, #15140122). Cells were washed with DPBS (Gibco, #14190144) and detached with 0.05% Trypsin-EDTA (Gibco, #25300054) before passaging onto imaging plates.

### Drug treatment

Nocodazole (MedChemExpress, #HY-13520) was treated at 5 μM for 0, 30, and 60-minutes for the three conditions, respectively. For the 30- and 60-minute conditions, nocodazole was also incubated during imaging.

### Fluorescent labeling

To label mitochondria, we used MitoTracker Green FM Dye (Invitrogen, #M46750) at 200 nM for 30 minutes. To label microtubule, we used 1X Tubulin Tracker Deep Red (Invitrogen, #T34076) for 30 minutes. During imaging, we also added 1X probenecid (Invitrogen, #P36400) to reduce the efflux of Tubulin Tracker Deep Red over the time of imaging.

### Lattice light sheet microscopy

A custom-built lattice light sheet microscope designed by Eric Betzig’s Lab at HHMI Janelia/UC Berkeley was used to image samples. Key modifications of this work are the use of 0.6 NA excitation objective lens (Thorlabs, TL20X-MPL), a 1.0 NA detection objective lens (Zeiss W Plan-Apochromat 20x/1.0, model #421452-9800), and a Hamamatsu Photonics Orca Fusion BT sCMOS Camera for image acquisition. 488nm and 560nm lasers were used to excite GFP and RFP. A Multiple Bessel Beam Light Sheet Pattern with NA Max 0.4, NA Min 0.35 was used for the organoid samples and a Hexagonal Lattice Light Sheet Pattern with NA Max 0.5 and NA Min 0.42 was used.

### Data preprocessing

Raw LLSM imaging data was preprocessed as described in Wang, Hakozaki, et. al (2024)^18^. Briefly, the LiveLattice preprocessing pipeline uses the WH-Transform, which combines deskewing, scaling, and rotation into a single memory-efficient transformation and allows real-time processing of data with a smaller data size.

### 4D segmentation

Individual Cal27 cells were segmented as follows. The fluorescence signal from MitoTracker Green was first normalized and smoothed. Membrane boundaries were estimated through spatial inclusion of the fluorescence signal where contours were created manually in 3D using IMOD^49^. The resulting 3D contours were converted to binary masks using a custom python script. Since cell movement was minimal over the duration of imaging (11 minutes), segmentation masks from the first frame were applied to all frames.

Nuclei for each cell were segmented in similar fashion to the cells where the location of the nuclear membrane was estimated based on the spatial exclusion of the MitoTracker fluorescence signal. 3D contours for both nuclear and membrane masks were then converted to binary masks using a custom python script.

Mitochondria segmentation was performed using MitoGraph^40^ (v3.0) with default parameters. The input to MitoGraph consisted of fluorescence signals from individual cells segmented previously. An adaptive thresholding approach with a block size of 10 pixels was used to compute a block-dependent local threshold. MitoGraph outputs include a segmented mitochondrial network along with a corresponding skeleton representation, which included network edges and node attributes such as 3D coordinates, fluorescence intensity, and tubular width. A custom Python script was used to process the MitoGraph outputs, reconstruct full-resolution mitochondrial networks, and compile the networks with node attributes into a python-igraph object.

Microtubules were segmented for each frame in 3D with the Allen Cell & Structure Segmenter^44^ plugin in Napari^45^ using default parameters for alpha tubulin. The resulting binary segmentation masks were then used as input for MitoGraph to obtain standardized outputs for tracking purposes and were additionally utilized separately for construction of microtubule models using custom python scripts.

### 4D organelle tracking

MitoTNT was utilized for quantifying both mitochondrial network and microtubule network dynamics^19^. Fragment-level (center-of-mass) motilities and remodeling rates were obtained from MitoTNT analyses of mitochondrial networks. In addition, node-level motilities were obtained from MitoTNT analyses of the segmented microtubule networks.

### Digital twin model construction

Custom factory methods were created for each digital twin model component derived from experimental data. These include the mitochondria, microtubules, membrane, and nucleus models. All models were constructed using only the first frame of 4D LLSM data. Python-igraph^50^ structures were used for defining all topological structures where spatial coordinates were assigned as node attributes. Note that voxel units (i.e., 3D pixel grid indices) are used for all coordinates and distances values unless explicitly stated otherwise. Voxel dimensions are 0.111 µm along x, y, and z dimensions and are reflective of the microscope’s resolution after data pre-processing. Functional descriptions for model construction routines are as follows.

Mitochondria network models were constructed using the spatially embedded python-igraph objects obtained from MitoGraph segmentation. Particle-based reconstructions of mitochondrial fragment topologies were generated by replacing nodes with particles and edges with harmonic potentials and positioned according to the node coordinate attribute. Mitochondria particles were assigned a radius of 0.15 µm as this both reflects fragment cross section radii observed experimentally^41^.

Plasma membrane models were constructed from their respective binary masks obtained from segmentation. To account for the reduction in volume due to non-finite particle sizes, cell masks were first dilated along all three using a custom 3D binary dilation method based on scikit-image’s binary dilation method^51^. Edge detection was then performed on the mask using a custom 3D implementation of canny edge detection based on scikit-image’s canny edge detection method^51^. Indices for non-zero pixels corresponding to the detected edge were collected and subsequently rescaled by the voxel dimensions to recover particle coordinates. Lastly, particle coordinates were down sampled using the farthest-point down sampling algorithm as implemented in Open3D^52^ to reduce the total number of particles in the simulation.

Nuclear membrane models were constructed from their respective binary masks obtained from segmentation. Edge detection was performed on the mask using a custom 3D implementation of canny edge detection based on scikit-image’s canny edge detection method^51^. Indices for non-zero pixels corresponding to the detected edge were collected and subsequently rescaled by the voxel dimensions to recover particle coordinates. To prevent overlap with other particle species (mitochondria or microtubules), particle coordinates were isometrically scaled about the center of mass.

Microtubule models were constructed from their respective binary masks obtained from segmentation. Binary segmentation masks for microtubules were first refined by intersecting the microtubule mask with the membrane and nucleus masks to ensure that all microtubule particles would reside within the cytosolic compartment without spatial overlap. Skeletonization was then performed in 3D using Lee’s algorithm implemented in scikit-image^51,53^. The resulting skeleton was converted into an undirected graph representation by assigning nodes for each pixel value where the pixel indices were assigned as a node attribute and edges were defined between spatially adjacent vertices within a predefined distance threshold. The initial graph was refined by removing edges connecting vertices with degrees exceeding a predefined degree threshold.

Disjoint graphs were then filtered based on a minimum number of nodes to remove noise segmentation artifacts. Regarding the reconstruction threshold values, a distance threshold of 2, maximum degree threshold of 3, and minimum node count threshold of 5 were found to produce good reconstructions with minimal artefacts.

The location of the centrosome was estimated using a custom algorithm employing a centroid-based intensity approach. With the centrosome typically proximal to the nuclear membrane, the nuclear membrane model was used to constrain the search space. Briefly, nucleus particle coordinates are iterated over and voxel indices spanning search regions are defined using a predefined search distance parameter. The raw microtubule fluorescence data is then sliced to isolate the corresponding search region, and a uniform threshold is applied using scikit-image^51^. If the candidate region size is greater than a predefined threshold and the total intensity exceeds that of the current region, the centrosome coordinate is updated using the indices of the maximum intensity voxel. We found that the centrosome location could be estimated reliably using a search distance threshold of 10 and centrosome cubic region volume threshold using a side length of 4.

Once identified, the centrosome was added as a node in the graph, and microtubules within a predefined distance were connected to the centrosome node. For the graph containing the newly defined centrosome node, shortest paths from each node to the centrosome were calculated using python-igraph’s built in method and subsequently used to redefine the edges to be retrograde-directed and to point towards the centrosome. For remaining disjoint subgraphs, polarity was randomly assigned through selecting a random terminal (degree=1) node as the root node and subsequently traversing the graph directing edges towards it. Finally, the entire graph was rescaled based on voxel dimensions to ensure proper physical representation in the simulation. To improve computational efficiency, the graph was down-sampled by collapsing n-consecutive internal (degree=2) nodes into a single node assigned the mean of the n coordinates.

### Reaction-diffusion simulation

*Compute resources*. All segmentation, tracking, and simulation tasks were performed on a desktop workstation with Ryzen 9 5950x 16-core processor and 128GB DDR4 RAM. All simulations were run on the CPU. Passive digital twin models required 15 minutes to simulate 2 minutes of cell time. Due to the increased number of particles and additional operations required for performing active transport, digital twin models incorporating active transport (including both control and nocodazole 30minute) required 2.5 hours to simulate 2 minutes of cell time.

*Particle representation and dynamics*. The reaction-diffusion simulations were implemented using ReaDDy, a particle-based reaction-diffusion simulation engine optimized for mesoscale biological modeling and trajectories were visualized using Simularium Viewer^42^. ReaDDy employs overdamped Langevin dynamics to simulate the motion of interacting particles in a spatially resolved environment. All reactions were evaluated according to a Kinetic Monte Carlo reaction scheme using ReaDDy’s implementation of the Gillespie stochastic simulation algorithm. For spatially dependent reactions, the sum of the reacting particles’ radii was used reaction radius *R*_*rxn*_.

The spatial domain was initialized with the 3D model components reconstructed from LLSM imaging data. All simulations were performed using non-periodic boundaries as defined by nuclear and plasma membrane particles which were assigned fixed positions. Mobile particle species (e.g., mitochondria, microtubules, motors) were contained by repulsive harmonic potentials defined between all sets of the membrane particles and all mobile particle species. This effectively constrained the simulation domain to the cytosolic compartment. Mitochondria were represented as collections of diffusing and interacting particles. Mitochondria topologies were reconstructed through replacing graph edges with harmonic bonds. Second-order harmonic angle potentials were introduced for mitochondria to reproduce experimental dynamics. The harmonic angle force constant was calibrated to reproduce mean-squared end-to-end distance for linear fragment graphs comprising 6 nodes sampled across all 60 frames from 4x cells treated from the 60-minute nocodazole treatment condition. Microtubules are similarly represented using particle topologies. Internal potentials defined for microtubule particles consisted of harmonic bonds and angles as topological and conformational constraints. Repulsive potentials were defined between microtubule particles to enforce spatial exclusion and for nuclear and plasma membranes to confine the network to the cytosolic compartment. Mitochondria and microtubule particles species experience stochastic motion governed by ReaDDy’s overdamped Langevin dynamics using a temperature of 37 C to match the temperature of the LLSM imaging chamber.

*Fusion-fission reactions*. Fusion and fission events were modeled as topological reactions and were parameterized using the results from MitoTNT analyses. From a kinetics perspective, fission is a spatially independent reaction because it requires only a single topology to occur where the observed or macroscopic fission rate is the product of the intrinsic or microscopic fission rate and the number of available fission sites. We therefore modeled fission events using a spatially independent structural topology reaction which randomly selects and removes a harmonic bond with a constant probability per bond per integration timestep to produce two new distinct fragments. Fusion requires an encounter between two distinct fragments which introduces spatial dependence. We therefore modeled mitochondrial fusion events using spatially dependent topological reactions where single fragment is formed by introducing a new harmonic bond is introduced between mitochondrial particles in the different fragments.

### Active transport modeling

We developed a decentralized active transport module for ReaDDy which enables simulation of motor-driven transport along network topologies. A spatially embedded directed graph structure is used to represent the microtubule network in the simulation where vertices and directed edges are mapped to particles and harmonic bonds. Motor proteins (e.g., dynein and kinesin) are modeled as two-state Markov models embedded as explicit particles which undergo stochastic state transitions and traverse the diffusing, polarized microtubule network according to their assigned polarity.

The probability of an event occurring within an integration timestep τ follows the general relation between microscopic reaction rates and discrete-time probabilities:

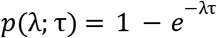

where *λ* and *p* represent the microscopic rate and the occurrence probability associated with a given event, respectively. This formulation is used for all stochastic events including motor stepping, activation, deactivation, binding, and unbinding. Motors are stepped according to a fixed velocity parameter along the microtubule contour if active. Inactive motors are stationary and maintain a constant relative position on the microtubule. Orientation-dependent binding was implemented for the first binding motor and subsequent motor types were selected with equal probability. We restricted mitochondria motor binding to terminal (degree=1) mitochondrial particles only where the number of motors was limited to one per terminal mitochondria particle. The number of motors bound to a single fragment topology was not limited. Thus, individual mitochondrial fragments may be simultaneously bound to multiple motor particles which leads to the possibility of a fragment being bound to and transported along one or more distinct microtubules.

Treating each microtubule particle as a binding site, newly created motors are assigned the position of the microtubule particle and take either an active state with probability *p* _*activate*_ or an inactive state with probability 1 − *p*_*activate*_. Each motor is registered with indices for the source and target vertices in the directed microtubule graph. The binding-site microtubule particle acts as the source vertex, while the target vertex is determined by the local directed edges and motor polarity. At each timestep, motors in any state may unbind with a probability *p*_*off*_ upon which they are immediately removed. Bound motors undergo stochastic transitions between active and inactive states according to the probabilities *p*_*activate*_ and *p*_*deactivate*_. Motor stepping is modeled as a stochastic process where each motor has a velocity *v* and step length *l*_*step*_ corresponding microscopic stepping rate is *λ*_*step*_ = *v*/*l* _*step*_. To enable multiple motor steps within a single timestep, motors step probabilities are re-parameterized to use a predefined global step length parameter *Ĩ*_*step*_ where step probabilities for each motor are adjusted to maintain a target velocity along the microtubule contour. When a motor has reached its target vertex, it is reassigned as the source and a new target vertex is selected based on the local directed edges and motor polarity. If a branching node (degree=3) is reached and multiple valid directed edges are available, the next target vertex was selected at random. To account for the diffusive motion of the underlying microtubule network, motor positions are corrected at each integration timestep to maintain accurate motor positions on the microtubule. Using the displacement vectors for source and target vertices across successive frames (δ_*source*_ and δ_*target*_), the motor position correction δ_*motor*_ is calculated as:

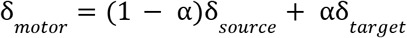

where α is the fractional position of the motor along the length of the edge. Given the motor step parameter *l*_*step*_ and the unit vector between source and target vertex positions, motor step position updates are calculated according to

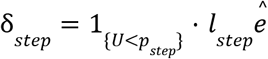

where 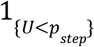 is an indicator function which operates on a random variable *U* sampled a uniform distribution *U*(0, 1) and takes a value of 1 if 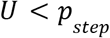 and 0 otherwise. The full motor is given by

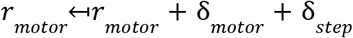

We additionally implemented force-dependent detachment whereby motors are removed if the motor-mitochondria bond energy exceeds a predefined threshold (**Table S3**). Motors are removed upon reaching the end of the microtubule which corresponds to a terminal (degree=1) node.

### Parameterization

Simulations for parameter calibration were performed in-twin using the cell models 60- and 0-minute condition in **Figure 1B** and **Figure 2B** unless explicitly stated otherwise. Diffusivities and remodeling event data were obtained from MitoTNT analysis results. Per-bond fission rates were calculated by dividing the number of events the frame delay followed by normalization by the number of mitochondrial nodes in the graph reconstructions. Fission rates were then evaluated according to the number of edges per-topology basis. The microscopic fission rate parameter was obtained by normalizing the macroscopic fission rates obtained from MitoTNT analyses by the edges in the corresponding network graphs for the 60-minute condition and were used for all simulations. The observed or macroscopic fusion rate such as that obtained from MitoTNT is a product of the intrinsic or microscopic rate (which lacks spatial dependence) and the frequency of such encounters (which incorporates spatial dependence). With experimentally observed fusion-fission rates being balanced such that the mean fragment size fluctuated stochastically about an equilibrium point, microscopic fusion reaction rates were iteratively tuned in-twin to maintain a mean fragment size over a period of two minutes. With the frequency of fragment encounters dependent on fragment motilities, fusion rate parameters for the 60- and control (0-minute) conditions were calibrated independently and in-twin using the digital twin models shown in **Figure 1B** and **Figure 2B**, respectively.

The diffusion coefficient for mitochondria particles was calibrated in-twin to match the 60-minute condition mean fragment diffusivity using simulations performed using the digital twin cell model shown in **Figure 1B** and used in all simulations. To calculate simulated mitochondrial fragment diffusivities, custom python scripts were used to parse trajectories to obtain per-topology trajectories, or a time-series of spatially embedded graph objects with a 1:1 correspondence to the particle topologies. Topology trajectory records are created for each initial fragment at t=0. Upon participating in either a fusion or fission reaction, the current topology trajectory is closed, and new trajectory records are initialized for each product topology. Each of the resulting topology trajectories was filtered so that only the desired species (i.e. mitochondria) was considered in the calculations.

To constrain the microtubule network to the cytosolic compartment, microtubule particles were assigned a radius of 0.05 µm, and repulsive potentials were defined between microtubules and both nuclear and plasma membrane species based on the sum of their respective radii. A mean edge length of 0.2 µm was used as the equilibrium bond length between microtubule particles. This was derived from the mean edge length in the down-sampled microtubule graph. We used a force constant of 0.05 kJ mol^-1^ nm^-2^ as we found it maintained constant distance between consecutive particles for the 5 ms integration timestep. Microtubule diffusivities were calibrated in-twin using the cell shown in **Figure 2B** to reproduce the mean node diffusivity obtained from MitoTNT analyses. Microtubule mechanical properties were modeled under the wormlike-chain model for polymer dynamics using second order harmonic angle potentials. Using an equilibrium angle of π rad, we calibrated the harmonic angle force constant using toy simulations to a persistence length of 1.4mm^54^ which was calculated via bond vector autocorrelation decay^55^.

Motor parameters were primarily drawn from published microkinetic values (**Table S3**). For the purposes of this study, we assumed active motors process at their maximum respective rates. The activation probability *p*_*activate*_ was the only free active transport parameter and was calibrated in-twin to reproduce the mean fragment-level diffusivity for the control condition with fusion-fission reactions enabled using the digital twin model shown in **Figure 2B**.

### Motility calculations

To quantify the motility of particles and filamentous fragments in our simulations, we computed the mean-squared-displacement (MSD) and extracted the corresponding diffusivity coefficients. The MSD for a given particle or fragment at time *t* is defined as

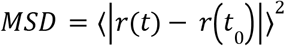

where *r*(*t*) represents a coordinate vector at time *t*. For each trajectory, particle positions are extracted from the simulation trajectory and are parsed to obtain per-particle trajectories unique particle identifiers. The positions were then used to construct an MSD matrix, where each column corresponds to a different particle or fragment, and each row represents a time point. Fragment level motility analyses were tracked according to the center-of-mass for each unique topology instance. Per-topology trajectories were extracted using the unique identities of the particles which comprise them. At each recorded frame, the fragment’s center of mass *R*(*t*) was computed from the positions of its constituent particles as

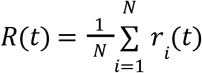

where *N* is the number of particles in the fragment and *r*_*i*_ (*t*) is the position vector for the *i*th particle in the topology at time *t*. The resulting center-of-mass coordinates were then used to construct an MSD matrix specific to fragments, allowing direct comparison between single-particle and fragment diffusion. The diffusivity *D* was obtained by fitting the MSD data to the linear relation

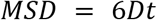

over an appropriate time window. We estimated the slope *m* of the MSD curve as

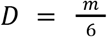

where *m* is determined from least-squares regression of the MSD values against their corresponding *t* values.

## Code Availability

The code used for constructing and performing all simulations is available at https://github.com/schoeneberglab/readdy-cell

## ACKNOWLEDGEMENTS

The authors thank Marta Medina and Gillian McMahon for helpful discussions. The authors thank Lisa Ilyin, Marta Medina, Dhruv Agarwal, and the other Schoeneberg Lab members for helpful suggestions during manuscript writing. This study was supported by grant 1R01GM148765-01 to JS.

## AUTHOR CONTRIBUTIONS

### Authors and affiliations

All authors were part of the Department of Pharmacology, University of California, San Diego, San Diego, CA, 92093 and the Department of Chemistry and Biochemistry, University of California, San Diego, San Diego, CA, 92093.

### Contributions

Eric Arkfeld and Johannes Schöneberg initiated the project. Eric Arkfeld performed 4D data processing, designed and implemented model construction pipelines and simulation framework, parameterized all models, and performed all simulations. Zichen Wang performed cell-culture, sample preparation for microscopy, and 4D data processing. Hiroyuki Hakozaki conducted LLSM Imaging. Eric Arkfeld and Johannes Schöneberg wrote the manuscript and were responsible for conceptualization. Johannes Schöneberg was responsible for funding and administration.

